# Quantification of Phase-Amplitude Coupling in Neuronal Oscillations: Comparison of Phase-Locking Value, Mean Vector Length, and Modulation Index

**DOI:** 10.1101/290361

**Authors:** Mareike J. Hülsemann, Dr. rer. nat, Ewald Naumann, Dr. rer. nat, Björn Rasch

## Abstract

Phase-amplitude coupling is a promising construct to study cognitive processes in electroencephalography (EEG) and magnetencephalography (MEG). Due to the novelty of the concept, various measures are used in the literature to calculate phase-amplitude coupling. Here, performance of the three most widely used phase-amplitude coupling measures – phase-locking value (PLV), mean vector length (MVL), and modulation index (MI) – is thoroughly compared with the help of simulated data. We combine advantages of previous reviews and use a realistic data simulation, examine moderators and provide inferential statistics for the comparison of all three indices of phase-amplitude coupling. Our analyses show that all three indices successfully differentiate coupling strength and coupling width when monophasic coupling is present. While the mean vector length was most sensitive to modulations in coupling strengths and width, biphasic coupling can solely be detected by the modulation index. Coupling values of all three indices were influenced by moderators including data length, signal-to-noise-ratio, and sampling rate when approaching Nyquist frequencies. The modulation index was most robust against confounding influences of these moderators. Based on our analyses, we recommend the modulation index for noisy and short data epochs with unknown forms of coupling. For high quality and long data epochs with monophasic coupling and a high signal-to-noise ratio, the use of the mean vector length is recommended. Ideally, both indices are reported simultaneously for one data set.

**Highlights:**

- mean vector length is most sensitive for differentiating coupling strength
- modulation index is most robust to differences in data length, sampling rate and SNR
- phase-locking value and mean vector length cannot detect biphasic phase-amplitude coupling

## 1. Introduction

Phase-amplitude coupling is a promising method to study cognitive processes (Jensen, 2006; Jensen and Lisman, 1998; Lisman and Jensen, 2013; Vosskuhl et al., 2015). There is no convention yet of how to calculate phase-amplitude coupling, but instead much heterogeneity of phase-amplitude calculation methods used in the literature. Most of these are reasonable measures from a theoretical point of view. To provide empirical evidence for choosing one of these measures over another, this work thoroughly compares the performance of the three most widely used phase-amplitude coupling measures with the help of simulated EEG data. The measures are the phase-locking value (PLV) by Mormann et al. (2005), mean vector length (MVL) by Canolty et al. (2006), and modulation index (MI) by Tort et al. (2008). From a historical viewpoint, the first amplitude modulations that have been detected are amplitude fluctuations of specific frequency bands, becoming apparent in the fast Fourier transform (FFT) of constituents of these signals (Burgess and Ali, 2002; Novak et al., 1992; Pfurtscheller, 1976). Because the FFT approach can solely reveal that the amplitude of a higher frequency oscillates at a lower frequency (characteristic of one signal), these amplitude modulations should not be misinterpreted to account for true temporal coupling between the instantaneous phase of the lower frequency and the amplitude envelope of the higher frequency (association between two signals and definition of phase-amplitude coupling). Neither the lower frequency itself nor its instantaneous phase are extracted in this approach.

Some of the most widely used phase-amplitude coupling measures today are the phase-locking value [PLV] (Mormann et al., 2005), also called synchronization index [SI] by Cohen (2008), the mean vector length [MVL] (Canolty et al., 2006), the modulation index [MI] (Tort et al., 2008), the envelope-to-signal correlation [ESC] (Bruns and Eckhorn, 2004), the general linear model approach [GLM] (Kramer and Eden, 2013; Penny et al., 2008), phase binning combined with analysis of variance (ANOVA) [BA] (Lakatos et al., 2005), and the weighted phase locking factor [wPLF] (Maris et al., 2011). All of these measures use the instantaneous phase and amplitude of band-pass filtered signals to calculate a measure that represents coupling strength. However, conceptual ideas and mathematical principles differ substantially between measures.

Several of these phase-amplitude coupling measures were compared with the help of simulated and real data in four reviews. Tort et al. (2010) executed the most extensive comparison so far, including most of the above listed measures and evaluating their performance pertaining to tolerance to noise, amplitude independence (independence from the amplitude of the amplitude-providing frequency band), sensitivity to multimodality, and sensitivity to modulation width. The modulation index, introduced by the same group (Tort et al., 2008), is well-rated in all aspects while, amongst others, the phase-locking value has poor ratings in all aspects. The mean vector length has good ratings in some aspects (e. g. tolerance to noise), but weaknesses in others (e. g. amplitude dependence).

Penny et al. (2008) introduced the GLM approach and compared it to the phase-locking value, mean vector length, and envelope-to-signal correlation in respect to noise level, coupling phase, data length, sample rate, signal non-stationarity, and multimodality. They found that the methods discriminated between data simulated with and without coupling to different extents, ranging from below chance level to perfect discrimination. Performance of the measures differed under poor conditions (high noise, low sampling rate, etc.), however, all measures performed equally well under good conditions (longer epochs, less noise, etc.).

Kramer and Eden (2013) introduced a new GLM cross-frequency coupling measure. It proves to be valid and performs equally well as the modulation index. The advantages of this method are that it can be interpreted as percentage change in amplitude strength due to modulation. Additionally confidence intervals are easily computed and the measure can detect biphasic coupling.

When Onslow et al. (2011), compared three phase-amplitude coupling measures (mean vector length, modulation index, cross-frequency coherence), they found that “no one measure unfailingly out-performed the others” (Onslow et al., 2011, p. 56). They concluded that each measure seems to be particularly suited for specific data conditions. Mean vector length for example is suitable for noisy data, exploratory analyses (analysing a broad frequency spectrum) and when the power of the amplitude providing frequency band is low.

The above cited reviews do not point to a single optimal measure for calculating phase-amplitude coupling. They rather show that most – but not all – of the used measures perform well and are equally affected by various confounders. Despite the availability of manifold measures, 79 % of studies use the phase-locking value adapted for phase-amplitude coupling, mean vector length, or modulation index (Hülsemann, 2016). Why is this the case? The phase-locking value is derived from a long-used, phase-phase coupling measure that is easily adapted for the purpose of phase-amplitude measurement. Its familiarity in the scientific community might have promoted its application. Possibly the predominant application of mean vector length is due to its mathematical directness. The modulation index is conceptually intuitive.

The majority of reviews used very straightforward data simulation methods. Oftentimes, a sinusoidal oscillation is constructed at a lower phase-providing frequency and at a higher amplitude-providing frequency. Phase-amplitude coupling is introduced by multiplying both signals (cf. Onslow et al., 2011, 1. p. 52). Amplitude is then extracted from the so constructed signal and phase is extracted from the pure sinusoidal oscillation of the lower frequency. White noise is added to both signals. There are two pitfalls in this approach. Both sinusoidal signals reflect a plain prototype of phase-amplitude coupling, but in real neuronal data, pure sinusoidal oscillation cannot be filtered; rather, frequency bands containing different amounts of various frequencies are extracted. Second, white noise is added to the simulated data, even though it is known that not white noise but Brownian noise is inherent to brain dynamics (He et al., 2010; Miller et al., 2009).

Because none of the hitherto existing reviews simultaneously meet the requirements of realistic simulation of EEG data, providing inferential statistics for comparison of the measures, investigating moderators of phase-amplitude coupling, and including the three most widely used measures (phase-locking value, mean vector length, and modulation index), a new comparison of these methods is presented here. We aim to combine the best aspects of all previous reviews. EEG data is simulated rather realistically according to the procedure described by Kramer and Eden (2013). The influence of several moderators (multimodality, data length, sampling rate, noise level, modulation strength, and modulation width) inspired by Tort et al. (2010) is investigated. Sensitivity and specificity of the phase-amplitude coupling measures are checked according to the methods described in Onslow et al. (2011). For all these comparisons, inferential statistics are provided.

## 2. Material and Methods

### 2.1. Simulation of EEG Data and Implementation of Phase-Amplitude Coupling

A characteristic of natural EEG data is the proportionality of its frequency spectrum to a power law P(f) ∼ (1/f^β^). Namely, the higher the frequency f, the weaker the amplitude P(f). The exponent β defines the strength of the amplitude decrease. White noise is defined by β = 0, pink noise by β = 1 and Brownian (red) noise by β = 2. Different investigations have shown that the frequency spectrum of human brain activity relates to Brownian (red) noise, with 2 < β < 3 (He et al., 2010; Miller et al., 2009). Because of this, Brownian noise was generated using MATLAB code provided by Zhivomirov (2013), in order to simulate EEG data (Figure 1A).

**Figure 1.**
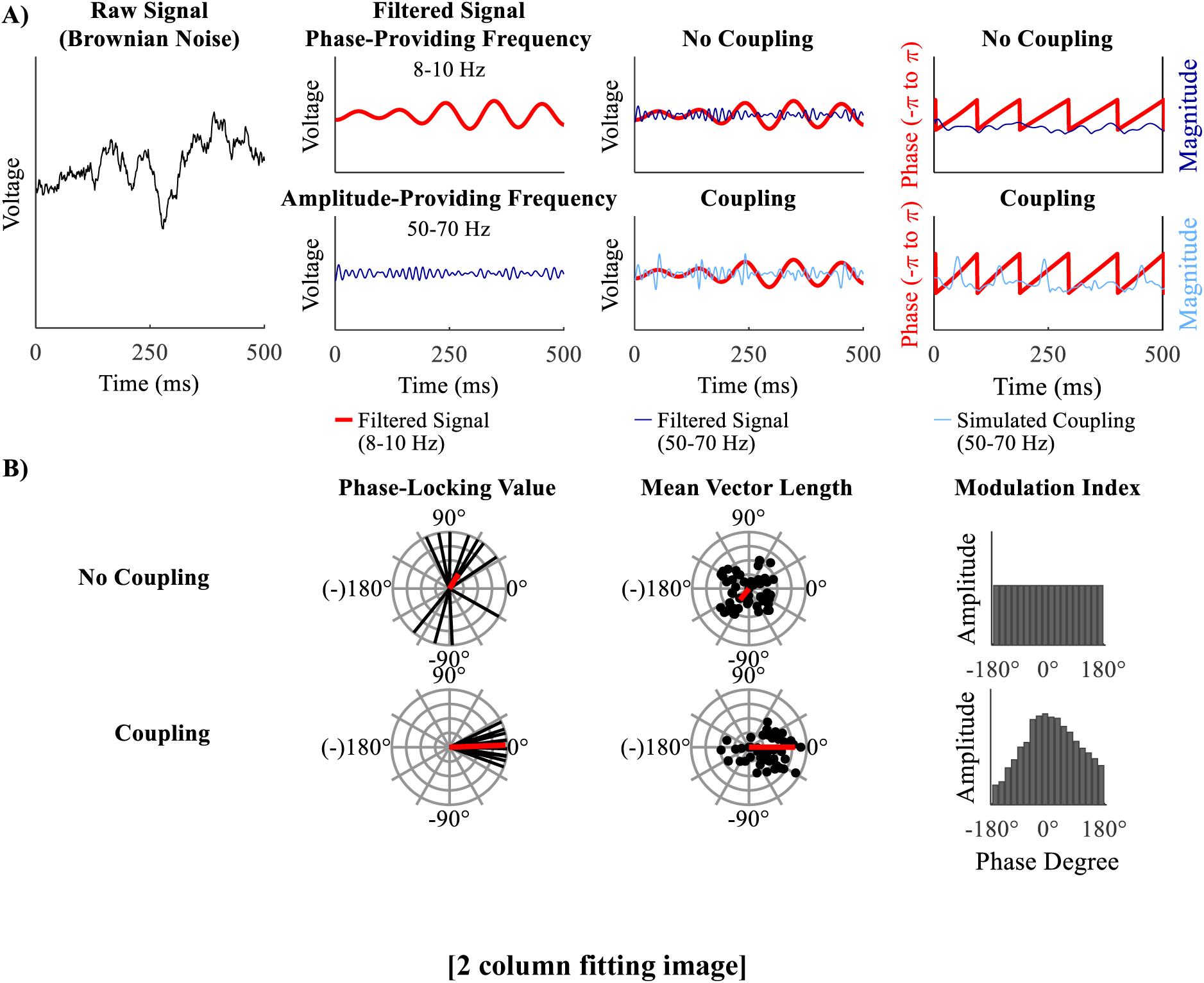
Simulation of the EEG signal and calculation of phase-amplitude coupling: A) (from left to right) Brownian noise is generated. This signal is band pass filtered to extract the slow phase-providing frequency (here 8-10 Hz, red line) and the fast amplitude-providing frequency (here 50-70 Hz, dark blue line). To simulate coupling (light blue line) the amplitude-providing band pass filtered signal is multiplied with a Hanning window plus one (not depicted here), which results in stronger amplitude at the peaks of the phase-providing frequency (lower middle right panel). Before extracting phase and amplitude (most right panels) band pass filtered noise (same frequencies) is added to the filtered data (not depicted here). The simulated coupling (light blue line) amplitude is most pronounced for phases at 0°. This is not the case for the original signal (dark blue line). B) Idealized depiction phase-locking value (left panels), mean vector length (middle panels), and modulation index (right panels) for a uniform distribution (upper panels) and phase-amplitude coupling (lower panels). Phase-Locking Value: Each black line represents the phase lag between two signals at one time point. The red vector is the mean of all black vectors. The upper panel shows inconsistent, widespread phase lags. The widespread phase lags lead to a relatively short mean vector (red line). The left panel shows an example of a relative constant phase lag around 0°. A relative constant phase lag leads to a relatively long mean vector (lower panel). Mean Vector Length: Each black dot represents one data point of the analytical signal. In case of coupling, a portion of the dots (or vectors) are especially long (reflecting strong amplitudes) at a specific narrow range of phase angles (here 0° in the lower panel). The red vector is the mean of all black vectors. It reflects coupling strength (short for no coupling – long for coupling). In case of phase-amplitude coupling it is indicating the preferred phase. Modulation Index: All possible phases are binned into 18 bins of 20° from -180° to 180°. Each bar reflects the mean amplitude of the amplitude-providing signal for the specified phase of the phase-providing frequency. This phase-amplitude plot is quantified with Shannon entropy. Shannon entropy is maximal for uniform distributions (upper panel). The Kullback-Leibler distance measures how much a given distribution (for example the one in the lower panel) deviates from the uniform distribution (depicted in the upper panel). The more phase-amplitude coupling there is in the data, the more the given phase-amplitude plot deviates from the uniform distribution and the higher the modulation index becomes.

Simulated data was then filtered at a low phase-providing frequency, from here on referred to as phase time series, with a narrow bandwidth of 2 Hz. The same data was filtered at a high amplitude-providing frequency, from here on referred to as amplitude time series, with a broad bandwidth. The exact bandwidth of the amplitude time series should depend on the frequency of the phase time series (Berman et al., 2012; Dvorak and Fenton, 2014). Because of this data was filtered, such that the sidebands of the modulating frequency were always included (i. e. centre frequency of amplitude-providing frequency band ± upper boundary of phase-providing frequency band).

A zero-phase Hamming-windowed sinc finite impulse response (FIR) filter implemented in EEGLAB (pop_eegfiltnew.m) was used. This function automatically chooses the optimal filter order and transition band width for a precisely selectable filter bandwidth. Low frequency was set to 8 – 10 Hz and high frequency to 50 – 70 Hz. Filtering can seriously distort raw data (Widmann et al., 2015), therefore only continuous data was filtered and first and last samples, where edge artefacts can occur, were later on discarded.

To introduce coupling, the procedure of Kramer and Eden (2013) was followed. A Hanning window plus one (i.e. each data point of the Hanning window is added with one) was multiplied with the amplitude time series. This multiplication of the Hanning window with the amplitude time series was not done continuously, but centred at either the relative maxima (peaks) or the relative maxima and minima (peaks and troughs) of the phase time series, in order to simulate monophasic and biphasic coupling, respectively. Extremum times are chosen because they are easy to detect. They relate to phase angles of 0° and 180°/-180°. Phase-amplitude coupling measures would not change if the coupling were to be introduced at another phase angle. The Hanning window itself is multiplied with the factor I to graduate the intensity of phase-amplitude coupling. To double the amplitude of the time series at the specified time I = 1.0 is chosen. I = 0.0 reflects no phase-amplitude coupling (i.e. not modulating the amplitude time series). The length of the Hanning window was also modulated to simulate different “widths” of phase-amplitude modulation. Parameters chosen for these moderators are specified below. In a final step, additional noise was added to the phase and amplitude time series. Therefore, Brownian noise of the same length was simulated, band-pass filtered at the same frequencies as the phase and amplitude time series, and added to the original phase and modulated amplitude time series, respectively. Frequency matched noise is disruptive to the modulated phase-amplitude coupling and therefore allows to check for the robustness of the phase-amplitude coupling measures.

Subsequently, phase and amplitude were extracted from the correspondent time series via Hilbert transform, using the Signal Processing Toolbox of MATLAB (The MathWorks, Inc). Then continuous phase and amplitude time series were segmented. This was done to introduce data discontinuities, which are present in real data as well. Filtering, Hilbert transform, and phase or amplitude extraction were always conducted on continuous data, to prevent filtering or other artefacts in the later analysed data epochs.

Data sets with a length of 42, 105, and 180 seconds were simulated. This amount of data is sufficient to simulate 30 trials with a length of 400, 2500 and 5000 milliseconds plus additional 30 seconds to introduce data discontinuities when segmenting the data. These parameters were chosen to mirror typical properties of event-related EEG data: (1) at least 30 trials per unique condition for which phase-amplitude coupling will be calculated (Luck, 2014), (2) trial length between 400 and 5000 milliseconds, and (3) data discontinuities between trials. Sampling rate was set to 1000 Hz (Cohen, 2014). In addition, simulated data was resampled to 500 Hz in order to investigate the influence of sampling rate. Noise was scaled by the factor 0.9, 1.0, and 1.1 in order to simulate different signal-to-noise ratios. Scaling factor 0.9, 1.0, and 1.1 correspond to a noise signal strength of 90 %, 100 %, and 110 % compared to the data signal strength. Four modulation strengths were realised: I = 0.0 for no coupling and I = 0.9, I = 1.0, and I = 1.1 for increasing coupling strength (I = 1.0 doubling the original amplitude strength).

These values lie within the range of former studies (e. g. Kramer and Eden, 2013). The length of the Hanning Window ranged between 22.5 % and 27.5 % of one low frequency cycle to modulate different “widths” of phase-amplitude modulation. This width is equivalent to about a quarter of one cycle and therefore covers the peak (or trough) phases of that low frequency cycle. At these phases, amplitude of the higher frequency was increased. All parameters were realised for mono- and biphasic coupling (factor multimodality).

### 2.2. Measuring Phase-Amplitude Coupling

To calculate phase-amplitude coupling, first, raw data is band-pass filtered in the frequency bands of interest. Second, the real-valued band-pass filtered signal is transformed into a complex-valued analytic signal. Finally, phase or amplitude is extracted from the complex-valued analytic signal. All these steps can essentially be implemented in MATLAB with four lines of code:

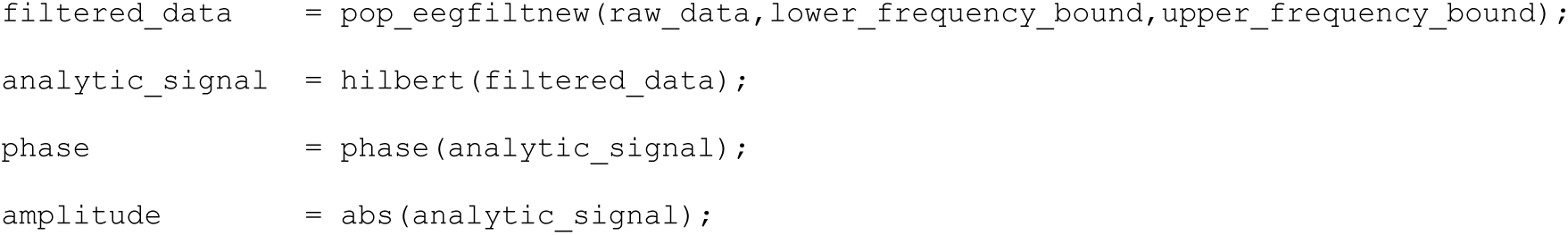

#### 2.2.1. Phase-Locking-Value by Mormann et al. (2005)

For the calculation of the phase-locking value, phase is extracted from the low frequency filtered analytic signal and amplitude is extracted from the high frequency filtered analytic signal. The amplitude time series is then again Hilbert transformed and phase is extracted from the “second” analytic signal. By these steps, one obtains phase angles for both time series for each data (time) point. For each time point the phase angle of the Hilbert transformed amplitude time series is subtracted from the phase angle of the phase time series, obtaining phase angle differences.

These phase angle differences can be plotted in a polar plane as vectors of the length one with the angle, representing the respective phase angle difference (Figure 1B, left panels). A constant phase lag between both time series indicates phase-amplitude coupling. A constant phase lag leads to vectors in the polar plane with a similar direction. Then all vectors are averaged: if they have a constant phase lag, they point into the same direction leading to a rather long mean vector. If there is a variable phase lag, the vectors are scattered around the polar plane, leading to a rather short mean vector. The length of the mean vector indicates the amount of phase-amplitude coupling (coupling strength). The direction of the vector represents the mean phase lag present between the two time series and the preferred coupling phase can be inferred from the phase lag. The phase-locking value is calculated by the following formula:

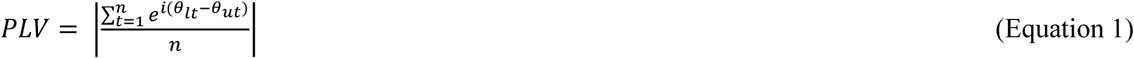

where n is the total number of data points, t is a data point, θ_lt_ is the phase angle of the lower frequency band at time point t and θ_ut_ is the phase angle of the Hilbert transformed upper frequency band amplitude time series.

The logic for this measure is that if and only if the amplitude of the high frequency time series oscillates at the lower frequency (indicator for phase-amplitude coupling) extracting instantaneous phase information from this signal will return valid phase angles that may have a constant phase lag to the instantaneous phase information of the low frequency band. If the amplitude of the high frequency time series does not oscillate at the lower frequency band (indicator for lack of phase-amplitude coupling), distorted phase information will be extracted from Hilbert transformed amplitude time series that will have an inconsistent phase lag to the instantaneous phase of the lower frequency signal.

One should be aware, that meaningful phase information can only be extracted from narrow band oscillations. The Hilbert transformed amplitude time series does not necessarily need to be such a narrow band oscillation.

#### 2.2.2. Mean Vector Length by Canolty et al. (2006)

The phase-amplitude coupling measure mean vector length (MVL) introduced by Canolty et al. (2006) utilizes phase angle and magnitude of each complex number (i. e. each data point) of the analytic signal in a quite direct way to estimate the degree of coupling. Each complex value of the analytic time series is a vector in the polar plane. Phase-amplitude coupling is present, when the magnitude M of a fraction of all vectors is especially high at a specific phase or at a narrow range of phases (Figure 1B, middle panels). Averaging all vectors creates a mean vector with a specific phase and length (red vector in Figure 1B). The length of this vector represents the amount of phase-amplitude coupling. The direction ∑ represents the mean phase where amplitude is strongest. When no coupling is present, all vectors cancel each other out and the mean vector will be short. Then its direction does not represent any meaningful phase. The mean vector length is calculated by the following formula:

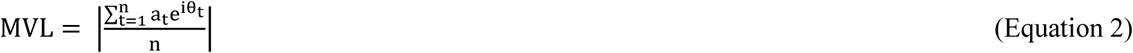

where n is the total number of data points, t is a data point, a_t_ is the amplitude at time point t and θ_t_ is the phase angle at time point t. This value cannot become negative because it represents the length of the mean vector. The length of a vector cannot be negative.

Three caveats come along with this measure: (1) the value is dependent on the general absolute amplitude of the amplitude providing frequency (independent of outliers), (2) amplitude outliers can strongly influence the mean vector length, and (3) phase angles are often not uniformly distributed (Cohen, 2014). All caveats are simultaneously counteracted by nonparametric permutation testing (see section 2.2.4). One of the reviews cited in the introduction (Tort et al., 2010) finds faults with the mean vector length being amplitude dependent. However, this is only true for the raw, but not for the permuted mean vector length.

In the interest of completeness, it should be mentioned that Özkurt and Schnitzler (2011) proposed a direct mean vector length which is amplitude-normalized and ranges between 0 and 1. When applying permutation testing to both mean vector length and direct mean vector length return essentially the same values. That is, when applied along with permutation testing, both measures are exchangeable. Without permutation testing, the usage of the direct mean vector length is recommended because it takes care of the possible amplitude differences in raw data.

#### 2.2.3. Modulation Index by Tort et al. (2008)

Tort et al. (2008) suggests a very different way of computing phase-amplitude coupling, which anyways is based on the same parameters of the analytic signal, amplitude magnitude and phase angle. For calculating the modulation index (MI) according to Tort et al. (2008), all possible phases from -180° to 180° are first binned into a freely chosen amount of bins. Tort et al. (2008) established to use 18 bins of 20° each, which many authors follow. The amount of bins can influence the results, as will be explaine x below. The average amplitude of the amplitude-providing frequency in each phase bin of the phase-providing frequency is computed and normalized by the following formula:

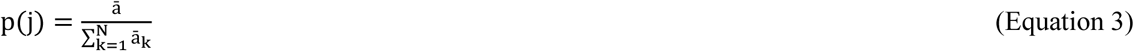

where ā is the average amplitude of one bin, k is the running index for the bins, and N is the total amount of bins; p is a vector of N values. With the help of these calculations, one obtains the data for the phase-amplitude plot, which depicts the actual phase-amplitude coupling graphically (Figure 1B, right panels). Subsequently Shannon entropy is computed; a measure that represents the inherent amount of information of a variable. If Shannon entropy is not maximal, there is redundancy and predictability in the variable. Shannon entropy is maximal, if the amplitude in each phase bin is equal (uniform distribution, Figure 1B, right upper panel). Shannon entropy is computed by the following formula:

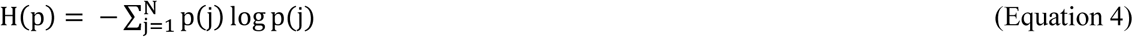

where p is the vector of normalized averaged amplitudes per phase bin and N is the total amount of bins. It does not matter which logarithm base is used if permutation testing is applied later on (Cohen, 2014). Like in Tort et al. (2008) the natural logarithm is used here. Shannon entropy is dependent on the amount of bins used and this is why the modulation index is likewise dependent on the number of bins. The higher the amount of bins, the larger Shannon entropy can become. Complying with the original author and most other studies, 18 bins have been employed here.

Phase-amplitude coupling is defined by a distribution that significantly deviates from the uniform distribution. Kullback-Leibler distance, a measure for the disparity of two distributions is calculated by the following formula:

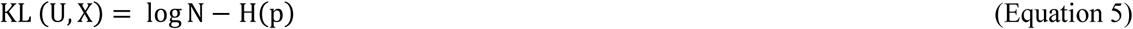

where U is the uniform distribution, X is the distribution of the data, N is the total amount of bins, and H(p) is the Shannon entropy according to equation 4. The uniform distribution is represented by log(N).

The final raw modulation index is calculated by the following formula:

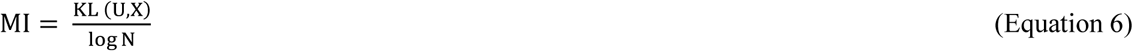

where KL(U,X) is the Kullback-Leibler distance according to equation 5 and N is the total amount of bins.

#### 2.2.4. Permutation Testing

All methods are subjected to permutation testing in order to quantify the meaningfulness of the derived value (Cohen, 2014). For permutation testing, the observed coupling value is compared to a distribution of shuffled coupling values. Shuffled coupling values are constructed by calculating the coupling value between the original phase time series and a permuted amplitude time series (or vice versa). The permuted amplitude time series is constructed by cutting the amplitude time series at a random time point and reversing the order of both parts. Generating surrogate data this way is most conservative, because it leaves all characteristics of the EEG data intact, except the studied one, namely the temporal relationship between phase angle and amplitude magnitude. Shuffling is usually repeated 200 to 1000 times (here we used 1000). The observed coupling value is standardized to the distribution of the shuffled coupling values according to the following formula:

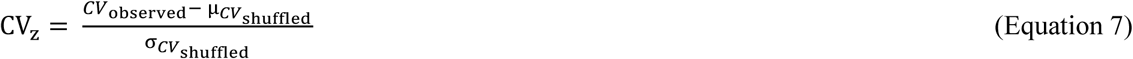

where CV denotes coupling value, μ denotes the mean and σ denotes the standard deviation (S. D.). Only when the observed phase-locking value is larger than 95 % of shuffled values (which are expected to be uncorrelated), it is defined as significant.

### 2.3. Statistical Analyses

All statistical analyses were conducted with IBM Statistics for Windows Version 23 (SPSS, Inc., IBM company), except otherwise specified. Significance level were set to *p* < .05. Violations of sphericity were, whenever appropriate corrected by Greenhouse-Geisser ε (Geisser and Greenhouse, 1958). Further analyses of significant results were conducted post hoc with Dunn’s multiple comparison procedure (Dunn, 1961) or post hoc t-tests. Effect size measure ω² is reported for significant results (Hays, 1973). It is an estimator for the population effect Ω², which specifies the systematic portion of variance in relation to the overall variance (Rasch et al., 2006).

#### 2.3.1. Specificity of phase-amplitude coupling measures

In a first step 10 000 data sets without coupling were simulated by setting the modulation strength to I = 0. Simulations were carried out for the frequency pair 8 – 10 Hz for phase time series and 50 – 70 Hz for amplitude time series. Phase-amplitude coupling values were generally compared in a 3 x 3 x 2 x 3 analysis of variance (ANOVA) with the repeated measurement factors method (phase-locking value, mean vector length, modulation index), data length (400 ms, 2500 ms, 5000 ms), sampling rate (500 Hz, 1000 Hz), and noise level (90 %, 100 %, 110 %).

As described above, nonparametric permutation testing was performed. Raw phase-amplitude coupling measures were z-standardized to the shuffled phase-amplitude coupling distribution. Normal z-values directly imply p-values; a value of 1.64 corresponds to a p-value of 5 %. The phase-amplitude coupling value distribution which is expected under the null-hypothesis does not have to match the standardised normal distribution. Therefore, significance was not inferred from the standardised normal distribution, but instead by that phase-amplitude coupling value, at which 5 % of simulated data (with no coupling) was classified as false positive. Shuffling for permutation testing was done within trials. Coupling measures were then calculated on concatenated trials.

Specificity of measures was analysed by counting false positives (significant coupling, even though it was not engineered into the simulated data) depending on (1) method, (2) data length, (3) sampling rate, and (4) noise level. To be able to conduct an ANOVA, the 10 000 simulations were divided into 100 subsamples of 100 simulations each. For each subsample false positives were counted. Each subsample was treated as a case in the subsequent 3 x 2 x 3 x 2 ANOVA with the repeated measurement factors method (phase-locking value, mean vector length, modulation index), data length (400 ms, 2500 ms, 5000 ms), sampling rate (500 Hz, 1000 Hz), and noise level (90 %, 100 %, 110 %) and the dependent variable false positives.

#### 2.3.2. Sensitivity of phase-amplitude coupling measures as a function of moderating variables

Performance of phase-amplitude coupling measures were quantified by simulating 100 independent data sets and modifying the parameters (1) modulation strength, and (2) modulation width, (3) multimodality, 1. (4) data length, (5) sampling rate, and (6) noise level within each dataset. Six 2-way ANOVAs were calculated. Each ANOVA included the repeated measurement factor method and was individually combined with the repeated measurement factors modulation strength (90 %, 100 %, 110 %), modulation width (22.5 %, 25.0 %, 27.5 % of one low frequency cycle), multimodality (monophasic, biphasic), data length (400 ms, 2500 ms, 5000 ms), sampling rate (500 Hz, 1000 Hz), and noise level (90 %, 100 %, 110 % compared to signal strength)

## 3. Results and Discussion

### 3.1. Specificity of Phase-Amplitude Coupling Measures

Phase-amplitude coupling values did not differ depending on method, data length, sampling rate, or noise level. Because of the high number of simulations (n = 10 000), some main effects and interactions became significant. However, all effect sizes were below ω² < .01, therefore these differences are negligible.

Figure 2 shows the phase-amplitude coupling value distribution for the phase-locking value, the mean vector length, and the modulation index. When setting the critical z-value for the phase-locking value at 1.86, for the mean vector length at 1.84, and for the modulation index at 1.92 five percent of the simulated data were classified as containing coupling (false positive). Thus, these values were defined as critical z-values. This implies that the mean vector length is most specific, directly followed by the phase-locking value. The modulation index is least specific compared to the two other methods.

**Figure 2.**
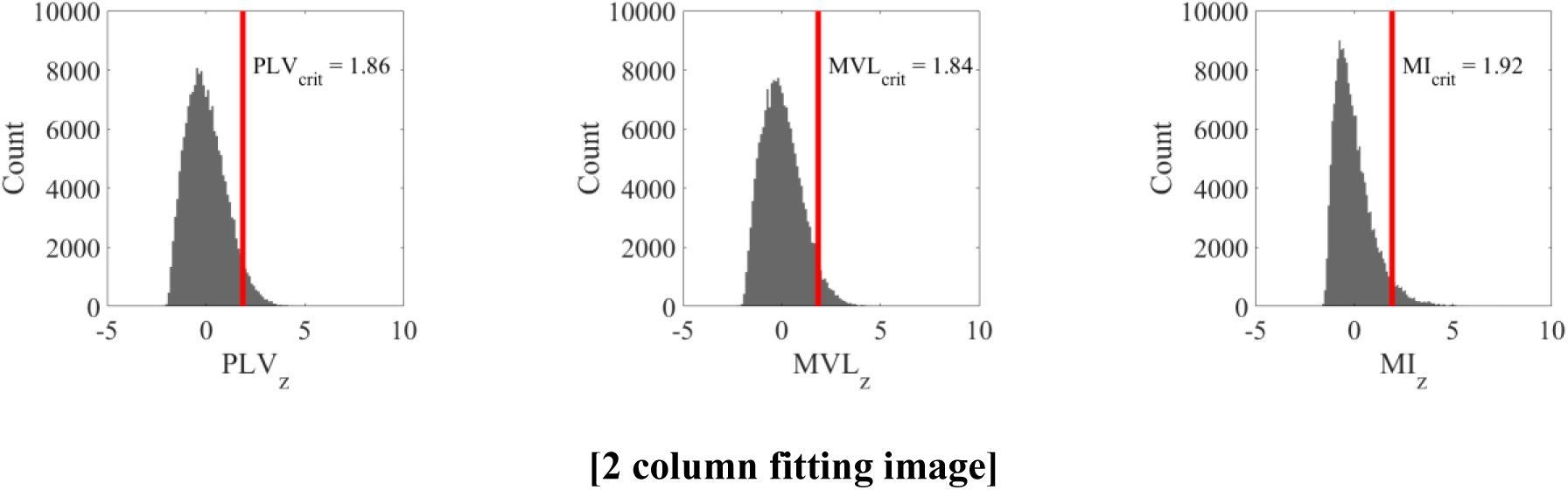
Probability distribution of coupling values under the null hypothesis: Phase-amplitude coupling value distribution under the null hypothesis (i. e. no coupling present in the data) of phase-locking value (left panel), mean vector length (centre panel), and modulation index (right panel). These distributions allow defining the significance threshold. The red line marks the critical phase-amplitude coupling z-value (relative cut off of 5 %). Choosing an absolute cut off instead would lead to smallest amount of false positives for mean vector length, followed by the phase-locking value. The modulation index would detect the most false positives.

The amount of false positives did differ depending on data length (*F*(2,198) = 27.19, *p* < .01, ω² = .15, Dunncrit = .26). There were significantly more false positives during short epochs (400 ms; mean ± S.E: 5.43 ± .07) compared to medium (2500 ms; mean ± S.E: 4.70 ± .06) and long epochs (5000 ms; mean ± S.E: 4.78 ± .09). Medium and long epochs did not differ in their false positive rates.

The main effect was qualified by a method by data length interaction (*F*(4,396) = 36.34, *p* < .01, ω² = .14, Dunn_crit_ = .23). This revealed that the above-described pattern was driven by the phase-locking value and mean vector length. There were no differences in false positive rate within the modulation index. Furthermore, in short epochs there were significantly more false positive in phase-locking value and mean vector length compared to the modulation index. In medium and long epochs there were significantly less false positive in phase-locking value and mean vector length compared to the modulation index.

Independently of the method, the main effect was further qualified by a sampling rate by data length interaction (*F*(2,198) = 36.14, *p* < .01, ω² = .10, Dunn_crit_ = .32). The above-described pattern of the main effect was only evident for a sampling rate of 500 Hz. Furthermore, the interaction revealed that there were more false positives for 500 Hz compared to 1000 Hz sampling rate during short epochs, no difference in false positives between sampling rates in medium epochs, and less false positives for 500 Hz compared to 1000 Hz sampling rate during long epochs.

### 3.2. Sensitivity of Phase-Amplitude Coupling Measures as a Function of Moderating Variables

#### 3.2.1. Effect of method on phase-amplitude coupling measures

Phase-locking value (1.66 ± .06) and mean vector length (2.08 ± .07) differed from the modulation index (11.97 ± .75) in their absolute magnitude independently of any other factor (main effect method: *F*(2,198) = 215.22, *p* < .01, ω² = .59, Dunn_crit_ = 1.34). Phase-locking value and mean vector length did not differ from each other.

#### 3.2.2. Effect of modulation strength on phase-amplitude coupling measures

Coupling values of all methods increased with increasing modulation strength (*F*(2,198) = 189.05, *p* < .01, ω² = .56). The interaction method by modulation strength became significant (*F*(4,396) = 151.54, p < .01, ω² = .40; Figure 3A). Post hoc t-tests showed that all factor levels within a method differed significantly from each other (all *p*’s < .01). The effect of modulation strength was most pronounced for the mean vector length (.47 < ω² < .76), followed by the modulation index (.49 < ω² < .69). The phase-locking value was least sensitive to modulation strength (.37 < ω² < .72).

**Figure 3.**
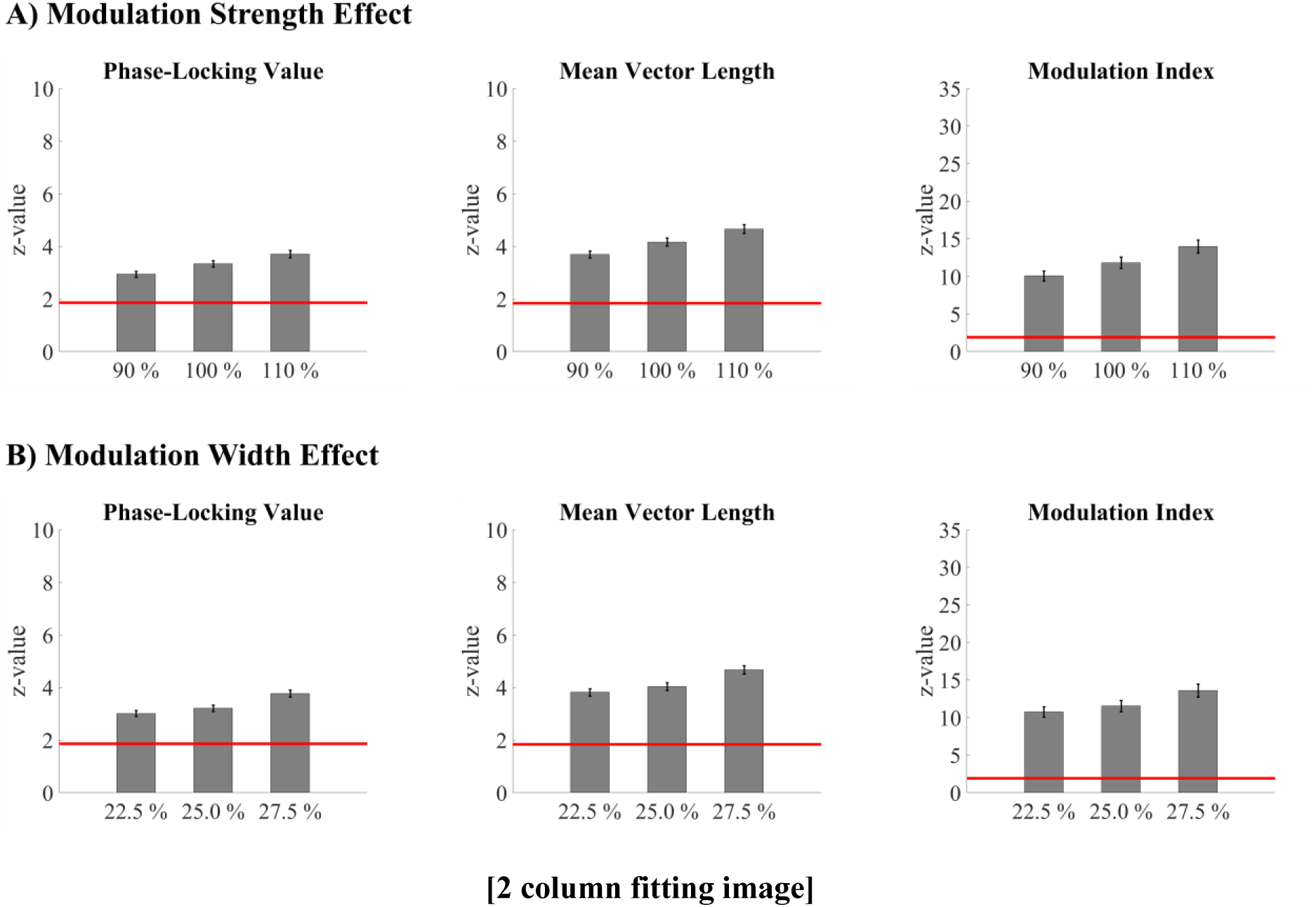
Sensitivity for modulation strength and width: Mean (± SEM) phase-amplitude coupling values for each method for the A) modulation strength effect and B) modulation width effect. Coupling values of all methods increased with increasing modulation strength. However, mean vector length differentiates best between the different factor levels of modulation strength. Also, coupling values of all methods increased with increasing modulation width. Here, phase-locking value and mean vector length differentiate best between the different factor levels of modulation width. The red line marks the significance level. All values above this line represent significant phase-amplitude coupling. For each effect, all factor levels within a method are significantly different from each other according to post-hoc t-tests. Only monophasic coupling values are depicted for the phase-locking value and the mean vector length.

The stronger the coupling, the larger phase-locking value, mean vector length, and modulation index are. As Tort et al. (2010) has shown, this behaviour is not inherent to all phase-amplitude coupling measures. Since researchers do not only want to prove the existence of phase-amplitude coupling, but also differentiate its strength, a measure that can do this is indispensable. Of all three methods, mean vector length differentiates best between the different factor levels of modulation strength.

#### 3.2.3. Effect of modulation width on phase-amplitude coupling measures

Coupling values of all methods increased with increasing modulation width (*F*(2,198) = 110.11, p < .01, ω² = .42). The interaction method by modulation width became significant (*F*(4,396) = 70.18, p < .01, ω² = .24; Figure 3B). Post hoc t-tests showed that all factor levels within a method differed significantly from each other (all *p*’s < .01). The effect of modulation width was most pronounced for the phase-locking value (.14 < ω² < .72) and the mean vector length (.14 < ω² < .71). The modulation index was least sensitive to modulation width (.15 < ω² < .52).

The broader the coupling width, the larger phase-locking value, mean vector length, and modulation index are. Of all three methods, phase-locking value and mean vector length differentiate best between the different factor levels of modulation width.

#### 3.2.4. Effect of multimodality on phase-amplitude coupling measures

Monophasic coupling (7.16 ± .36) led to overall stronger coupling measures than biphasic coupling (3.31 ± .24; *F*(1,99) = 813.94, *p* < .01, ω² = .80). Biphasic coupling could not be detected by the phase-locking value (3.33 ± .12 vs. -01. ± .01; *t*(99) = 27.26, *p* < .01, ω² = .88) and mean vector length (4.17 ± .15 vs. -01. ± .01; *t*(99) = 27,85, *p* < .01, ω² = .89). The modulation index was larger in monophasic than in biphasic coupling (13.98 vs. 9.96; *t*(1,99) = 22.49, *p* < .01, ω² = .83; Figure 4A).

**Figure 4.**
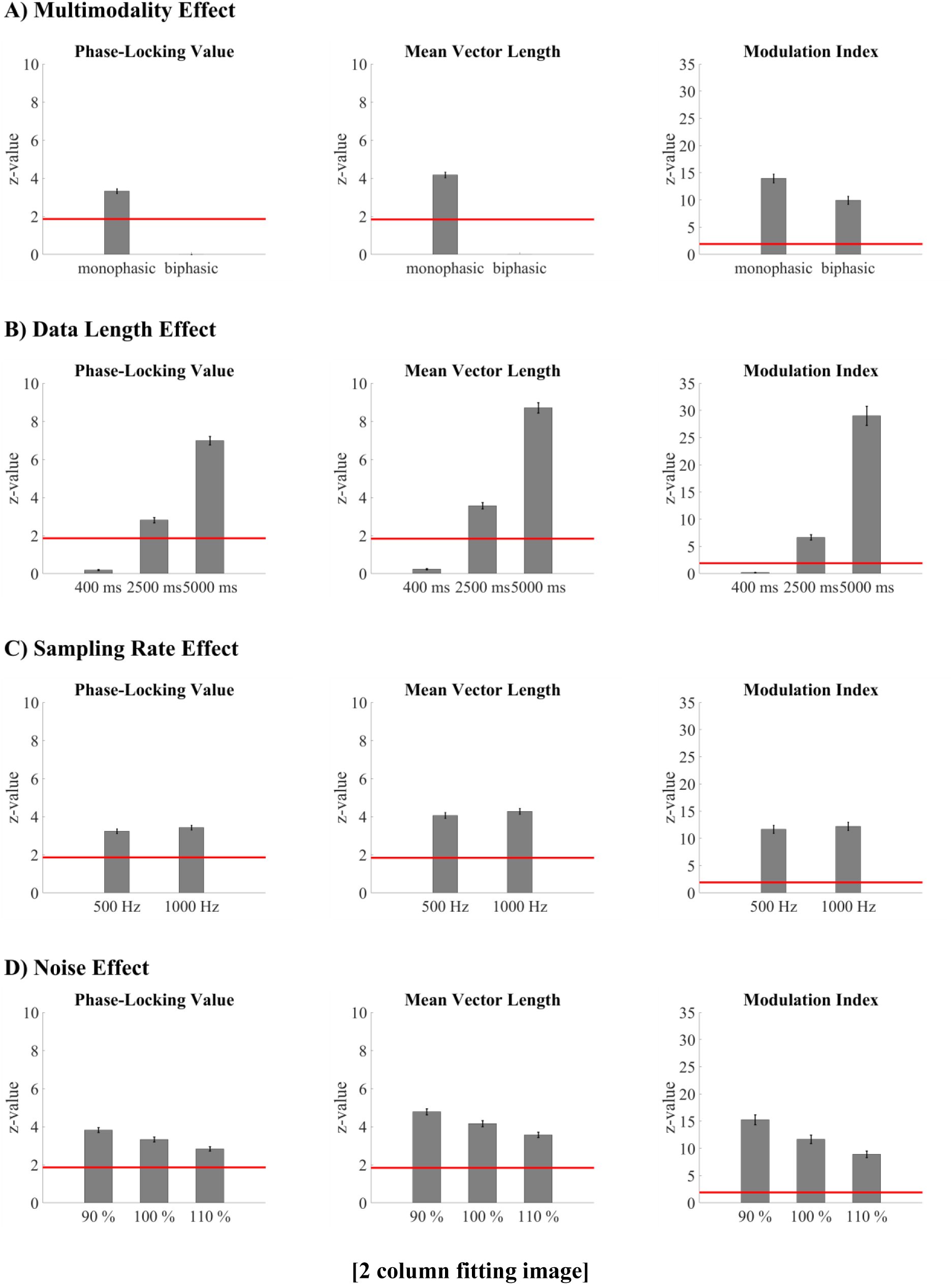
Moderators of the phase-amplitude coupling measures: Mean (± SEM) phase-amplitude coupling values for each method for the A) multimodality effect, B) data length effect, C) sampling rate effect, and D) noise effect. In contrast to the modulation index, biphasic coupling could not be detected by the phase-locking value and mean vector length. This factor might turn out to be not as important, as most studies report monophasic coupling. Coupling values of all methods increased with increasing data length and slightly increase with sampling rate. Sampling rate only becomes relevant when analysing frequencies close to the Nyquist frequency. Of all three methods, modulation index is least affected from the confounding factor data length and sampling rate. Coupling values of all methods decreased with increasing noise, while the phase-locking value is least affected from this confounding factor. The red line marks the significance level. All values above this line represent significant phase-amplitude coupling. For each effect, all factor levels within a method are significantly different from each other according to post-hoc t-tests. For B), C), and D) only monophasic coupling values are depicted for the phase-locking value and the mean vector length.

That is, multimodality influences the three methods very differently. Phase-locking value and mean vector length cannot find biphasic coupling as it was implemented here (amplitude of the higher frequency was increased at peak and trough of the lower frequency). Because of the mathematic construct of the mean vector length (equation 2, Figure 1B) this is not surprising. Peak and trough appear on opposite sides in the polar plane: their mean will cancel each other out. If other forms of biphasic coupling would be present, the mean vector length could be able to find it, but would probably underestimate its strength and would furthermore return distorted phase information. Therefore, it is important to have a look at the polar plot before interpreting one’s results. Similarly, the phase-locking value cannot detect biphasic coupling, as it was implemented here. For biphasic coupling the amplitude envelope oscillates twice as fast as the lower frequency band. Because of this, the phase lag between lower and upper frequency band spans the entire polar plane. The modulation index is able to find biphasic coupling, but biphasic coupling leads to a reduction in the phase-amplitude coupling value. Literature indicates that biphasic coupling plays a minor role in empiric data. To our knowledge only a very small fraction of studies report biphasic coupling (e. g. Lega et al., 2016, Leszczynski et al., 2015, van der Meij et al., 2012). Most studies report monophasic coupling.

#### 3.2.5. Effect of data length on phase-amplitude coupling measures

Coupling values of all methods increased with increasing data length (main effect data length: *F*(2,198) = 349.13, *p* < .01, ω² = .70). For the shortest epoch of 400 ms, none of the methods could detect significant coupling, even though it was engineered into the data. The interaction method by data length (*F*(4,396) = 240.65, *p* < .01, ω² = .52; Figure 4B) became significant. Post hoc t-tests showed that all factor levels within a method differed significantly from each other (all *p*’s < .01). The data length effect was most pronounced for mean vector length (.82 < ω² < .94), and phase-locking value (.80 < ω² < .94). The modulation index was least affected by data length (.65 < ω² < .75).

Overall, the longer the data, the larger phase-locking value, mean vector length, and modulation index are. This association was found in the data presented here, but must not generally apply. Here coupling was simulated continuously into the data. If coupling is transient and does not proportionally vary with data length, this relationship does not need to apply. Penny et al. (2008) showed, that coupling strength decreases for phase-amplitude coupling, which was simulated transiently. Potentially, the general rule is that the longer the data epochs where coupling occurs, the stronger the phase-amplitude coupling values. This should be tested in a follow-up analysis. This analysis further showed that a minimal data length is required for finding coupling, which should exceed at least 400 milliseconds per trial when including 30 trials (also see Cheng et al., 2018). None of the methods were able to detect coupling in the shortest simulated epoch of 400 milliseconds. It might be useful to develop a correction factor (e. g. similar to the pairwise phase consistency that is insensitive to data length variation; Vinck et al., 2010) for data length, to make phase-amplitude coupling values more comparable across studies. Of all three methods, modulation index is least affected from the confounding factor data length.

#### 3.2.6. Effect of sampling rate on phase-amplitude coupling measures

Overall coupling values slightly increased with increasing sampling rate (*F*(1,99) = 23.65, p < .01, ω² = .10). The sampling rate effect differed according to the method (*F*(2,198) = 14.02, *p* < .01, ω² = .04; Figure 4C). It was most pronounced in the mean vector length (*t*(99) = -5.15, *p* < .01, ω² = .20), followed by the phase-locking value (*t*(99) = -4.86, *p* < .01, ω² = .18). The modulation index was least affected by sampling rate (*t*(99) = -4.23, *p* < .01, ω² = .14).

The factor sampling rate stands out because of its comparatively small effect size. A second set of data was simulated testing phase-locking value, mean vector length, and modulation index at 16 – 18 Hz for the modulating frequency and 202 – 238 Hz for the modulated frequency (for detailed results see Hülsemann, 2016). This analysis showed that sampling rate is indeed important, but only if the investigated upper frequency band approaches the Nyquist frequency. Of all three methods, modulation index is least affected from the confounding factor sampling rate.

#### 3.2.7. Effect of noise on phase-amplitude coupling measures

Coupling values of all methods decreased with increasing noise (*F*(2,198) = 325.22, p < .01, ω² = .68). The interaction method by noise became significant (*F*(4,396) = 251.00, p < .01, ω² = .53; Figure 4D).

Post hoc t-tests showed that all factor levels within a method differed significantly from each other (all *p*’s < .01). The effect of noise was most pronounced for the modulation index (.65 < ω² < .76) and the mean vector length (.55 < ω² < .84). The phase-locking value was least affected by noise (.51 < ω² < .81).

Overall, the noisier the data, the lower phase-locking value, mean vector length, and modulation index are. This aspect is not desired but plausible. Noise obscures the relation between the phase of the lower frequency and amplitude of the higher frequency. The data as a whole contains phase-amplitude coupling to a lesser extent, as the relative amount of noise compared to the relative amount of signal increases. Of all three methods, phase-locking value is least affected from the confounding factor noise.

#### 3.2.8. Interaction Effects

Conducting a 6-way ANOVAs for each method separately (see Hülsemann, 2016 for detailed results), revealed ordinal interaction for all factors (multimodality, data length, sampling rate, noise, modulation strength, and modulation width). Especially multimodality and data length interacted with the remaining factors, as well as interacted with each other and the remaining factors. Sampling rate only showed significant interactions, when analysing frequencies close to the Nyquist frequency. All interactions had a monotone pattern, following the pattern of each main effect. For example, mean vector length increased the longer the data, but it increased less when also noise increases (Figure 5). This pattern was true for each added factor. Phase-locking value and mean vector length did not find biphasic coupling at all. Because of this, for these two methods, the described main effect and interaction patterns are only valid for monophasic, but not for biphasic coupling. For the modulation index the pattern was true for mono- and for biphasic coupling.

**Figure 5.**
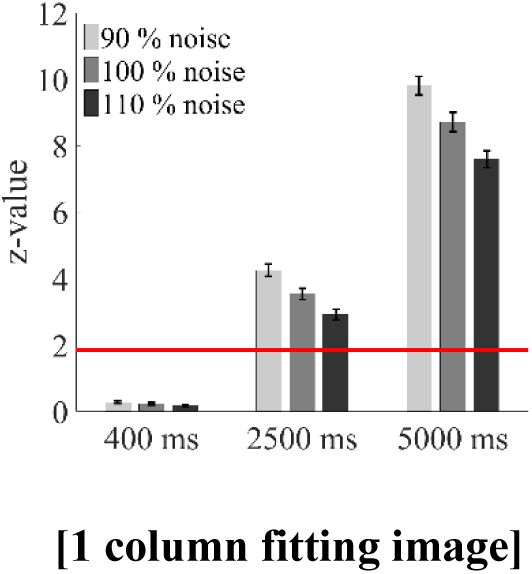
Interaction effects between the moderators of the phase-amplitude coupling measures: Mean (± SEM) phase-amplitude coupling values for the mean vector length for the data length by noise interaction (only monophasic coupling values). Interactions had a monotone pattern, following the pattern of each main effect. Depicted here, mean vector length increased the longer the data, but it increased less when also noise increased. This pattern was true for each added factor. The red line marks the significance level. All values above this line represent significant phase-amplitude coupling. For each method, all factor levels are significantly different from each other according to Dunn’s post hoc test. Only values within the 400 ms condition do not differ between the noise levels.

Comparing all three methods it becomes evident that the modulation index is least affected by the confounding factors multimodality, data length and sampling rate. However, it is also – like the phase-locking value – less sensitive to variation in modulations strength compared with the mean vector length. The modulation index is especially less sensitive to modulation width compared to the mean vector length and phase-locking value. Mean vector length and modulation index are similarly – and stronger than the phase-locking value – affected by the confounding factor noise.

## 4. Conclusion

In conclusion, for long data epochs, recorded at high sampling rates, with a high signal-to-noise ratio, the use of the mean vector length is recommended, because it is more sensitive to modulation strength and width than both other methods. For noisier data, shorter data epochs, recorded at a lower sampling rate, the use of the modulation index is recommended, as it is least influenced by the confounding factors compared with both other methods. If it is not clear whether cross-frequency coupling will be mono- or biphasic, the modulation index should be used, even though literature suggests that biphasic coupling can be neglected.

The phase-locking value does not stand out in comparison to the two other measures. Its usage is potentially problematic because phase information is extracted from the amplitude envelope of a signal. Phase information can only be correctly extracted from truly oscillating signals; this must not be necessarily the case for an amplitude envelope. So far, no review evaluated this measure explicitly as positive.

Because mean vector length and modulation index have complementing strengths and weaknesses, it would be advisably to calculate both. The time-consuming aspect of measuring phase-amplitude coupling is permutation testing. Calculation of both measures on the other hand will not substantially increase the analysis time.

The modulation index is quantitatively larger than the phase-locking value and mean vector length. However, even despite substantial quantitative differences in values, the qualitative decision for significance of phase-amplitude coupling is the same for all three methods in our simulation. Nevertheless, comparison of coupling strengths between the methods is problematic and this lack of comparability provides another reason for reporting both, mean vector length and modulation index.

In contrast to mean vector length, the false positive rate of the modulation index is not affected by any confounding factor. However, this advantage against mean vector length is counteracted by one disadvantage against the mean vector length: calculation of the modulation index includes Shannon’s Entropy. The entropy value depends on the amount of bins as well as amount of data squeezed into the same amount of bins. This is an undesirable degree of freedom, which is not present when calculating the mean vector length.

Due to the dependency on confounding variables (e. g. data length), comparing absolute coupling strengths across studies might be difficult even if using the same method. Comparisons within one study, on the other hand, can be done with confidence. Nevertheless, one should make sure that signal-to-noise ratio is comparable within all experimental conditions and over the course of the experiment.

Generally, it is advisable to work with standardized phase-amplitude coupling measures via permutation testing. It facilitates the interpretation of the measures, first and foremost, by giving the researcher knowledge about the probability that the observed modulation index would have been also found under the assumption of the null-hypothesis. This aspect is often ignored in the literature.

Kramer and Eden (2013) stated that “an optimal analysis method to assess this cross-frequency coupling (CFC) does not yet exist” (p.64). Even if it would be ideal, to have a measure that is less susceptible to confounding variables summarizing this analysis, it should be rather concluded that at least two reasonable analysis methods exist.

